# The role of sensorimotor variability and computation in elderly’s falls

**DOI:** 10.1101/196584

**Authors:** Chin-Hsuan Lin, A Aldo Faisal

## Abstract

The relationship between sensorimotor variability and falls in elderly has not been well investigated. We designed and used a motor task having shared biomechanics of walking and obstacle negotiation to quantify sensorimotor variability related to locomotion across age. We also applied sensory psychophysics to pinpoint specific sensory systems associated with sensorimotor variability. We found that sensorimotor variability in foot placement increases continuously with age. We further showed that increased sensory variability, specifically increased proprioceptive variability, the vital cause of more variable foot placement in the elderly. Notably, elderly participants relied more on the vision to judge their own foot’s height compared to the young, suggesting a shift in multisensory integration strategy to compensate for degenerated proprioception. We further modelled the probability of tripping-over based on the relationship between sensorimotor variability and age and found a good correspondence between model prediction and community-based data. We revealed increased sensorimotor variability, modulated by sensation precision, a potentially vital mechanism of raised tripping-over and thus fall events in the elderly. Therefore, our tasks, which quantify sensorimotor variability, can be used for trip-over probability assessment and, with adjustments, potentially applied as a training program to mitigate trip-over risk.

## Introduction

Falls are the leading cause of unintentional injuries in the elderly^1, 2^; they account for more than 60% unintentional injuries and 50% of accidental death among people aged 65 and older^1^. Most falls in the elderly occur during locomotion, and more than 50% of falls are triggered by tripping over something during walking^3–7^. Biomechanical research has documented gait differences between the young and elderly that are related to falls. Among them, gait variability has been consistently observed to increase with age^8–10^, and elderly fallers have higher variability than elderly non-fallers^8^. Increased gait variability is thus considered a marker of increased risk of falling. Although these epidemiological and biomechanical studies have provided phenomenological understandings of falls, the physiological basis underlying elderly’s gait variability changes remains largely unexplored.

Walking, like other motor tasks, is achieved by making use of sensory information, which conveys the states of the environment and body, to generate and execute motor commands^11–13^. The neural signals, from sensation, motor planning to execution, are intrinsically noisy^11, 14^. This noise in the sensory and motor systems causes variations of motor performance across multiple repetitions of a task, i.e. sensorimotor variability^13–15^. Most studies probing the ageing effects on sensorimotor tasks have focused on the upper limb and consistently found higher sensorimotor variability in the elderly^16–19^. This increase of sensorimotor variability is associated with a deficiency in elderly’s activities of daily living^16, 20^. A small number of studies compared ankle positioning precision between the young and elderly^21, 22^ and showed increased position variability in the elderly. It is thus reasonable to hypothesise that increased sensorimotor variability with age also presents in locomotion and manifests as increased gait variability. Further, this raised sensorimotor variability in the lower limb would impact negatively on elderly’s health and quality of life by increasing their fall risk.

Evidence has supported that ageing-related deficits of multiple sensorimotor functions are linked to gait decline and an increased probability of falling^23–27^. However, the majority of these studies focused only on average performance metrics, but not variability. One study compared quadriceps muscle force variability among young, elderly non-fallers and fallers^25^ and showed a link between greater force unsteadiness and fall history. Regarding sensation, the roles of vision and proprioception playing in falls have been investigated mainly through balance control. For example, studies have revealed that raised ankle joint position sensation variability during ageing are likely to contribute to impaired static postural control in the elderly^28, 29^. Much less is known about the association between sensory deficits during physiological ageing and sensorimotor variability, let alone gait variability. We argue that there is a need to delineate the connection between sensory variability and gait impairments with age because this approach links sensory degradation to trip-over risk when walking.

There is increasing evidence showing that people make use of visual estimates of the environment (see review^30^) as well as visual^30^ and proprioceptive estimates of the body position^31^ to guide their locomotion. The gross sensory variability of walking thus originates from the visual uncertainty of the environment as well as the visual and proprioceptive uncertainty of the body. Vision^23, 32^ and proprioception^31, 33^ are susceptible to ageing process, but the scales and speed of changes may differ. To date, there has been no systemic investigation to pinpoint the relevance of the variability of individual sensory modalities to ageing-related sensorimotor variability alterations. As modality specific measurements can lead to a more precise understanding of mechanisms behind increased sensorimotor and gait variability and thus the development of targeted fall prevention strategies, it is valuable to evaluate visual and proprioceptive variability separately.

Of many gait variables, minimum foot clearance (MFC), the minimum vertical distance between the lowest surface of the foot/shoe and the ground surface during the mid-swing phase of gait, is considered to be directly linked to trip-over occurrence^8, 10, 34, 35^. This is because that a foot or obstacle-ground encounter (i.e. a trip-over) occurs when MFC equals to or is lower than zero^8, 10^. Extensive investigations during the past two decades have identified greater MFC variability in the elderly, especially elderly fallers in multiple studies and regarded it as an important risk factor for falls^8, 10, 34, 35^.

Taking the evidence as mentioned above together, we hypothesised that increased sensorimotor variability with age manifests as increased MFC variability and thus increased trip-overs and falls. Moreover, raised sensory variability due to degeneration contributes to the increase of sensorimotor variability. The objective of this study is thus to characterise sensorimotor and sensory variability in the context of MFC. To do so, we designed and used a controlled motor task biomechanically mimicking stepping, FOot HEight POsitioning (FOHEPO) task, to compare sensorimotor variability in different age groups. We then used three sensory psychophysics experiments and sensorimotor computation models to quantify not only overall sensory variability but also visual and proprioceptive variability individually in context of the FOHEPO task. This approach enabled us to correlate sensory variability with sensorimotor variability, MFC variability and trip-over risk. We additionally assessed the relationship between sensorimotor variability and ages and performed modelling to indicate the probability of tripping-over while negotiating structured obstacles (i.e. staircases).

## Results

### Experiment 1: Sensorimotor variability of the FOHEPO task

We designed and applied the FOot HEight POsitioning (FOHEPO) task to measure end-point variability during movements that were biomechanically comparable to foot clearance when stepping to elevated surfaces. We used standard deviations to represent the sensorimotor variability of the FOHEPO task because Kolmogorov-Smirnov Goodness-of-Fit tests confirmed the normality of the foot height distributions (supplementary material Fig. 1). The mean sensorimotor variability measured for each group of each condition can be seen in Table 1. Sensorimotor variability increased significantly with age, as revealed by Age *×* Height *×* foot ANOVAs (age main effect: *F*_(2,70)_ = 4.71; *p* = .012). Post-hoc analysis displayed a significant difference between the elderly and young (*p* = .003), but not the middle-aged and young (*p* = .63). A significant height effect was also found (*F*_(2,140)_ = 55.13; *p < .*001). Further, because of a marginally significant age-group/obstacle height interaction (*F*_(4,140)_ = 2.04, *p* = .09), we analysed the simple effect of age on each height condition using Age *×* foot ANOVAs. Elderly performed significantly more variably than young when matching obstacles at 100 mm (*p* = .012) and 150 mm (*p* = .014) (see Fig. 3 a.). We performed a multiple linear regression, predicting sensorimotor variability from age, foot and obstacle height and found a significant positive correlation (*F*_(5;72)_ = 21 : 27; *p* < .001, adjusted *R*^2^ = .19) (black dots and line in Fig. 3 c.). All variables, except foot, added statistical significance to the prediction with *p* < .05. The result of *R*^2^ indicated that 19% of the variability in the data was explained by age and obstacle height. A test of quadratic trend was statistically non-significant (*p* = .20)

**Table 1.** Sensorimotor variability by age groups, feet and heights. Expressed in mean(SD) unit: mm

|  | Young | Middle-aged | Old |
| --- | --- | --- | --- |
| **L50** | 10.0(4.0) | 10.5(4.4) | 11.2(4.3) |
| **R50** | 8.6(2.7) | 11.3(5.0) | 11.3(5.2) |
| **L100** | 12.0(4.2) | 12.1(4.2) | 14.8(5.9) |
| **R100** | 10.7(3.7) | 12.6(5.5) | 14.8(5.5) |
| **L150** | 13.6(3.5) | 15.3(7.5) | 17.5(7.2) |
| **R150** | 12.2(3.8) | 14.4(6.8) | 17.2(7.3) |

**Figure 1.**
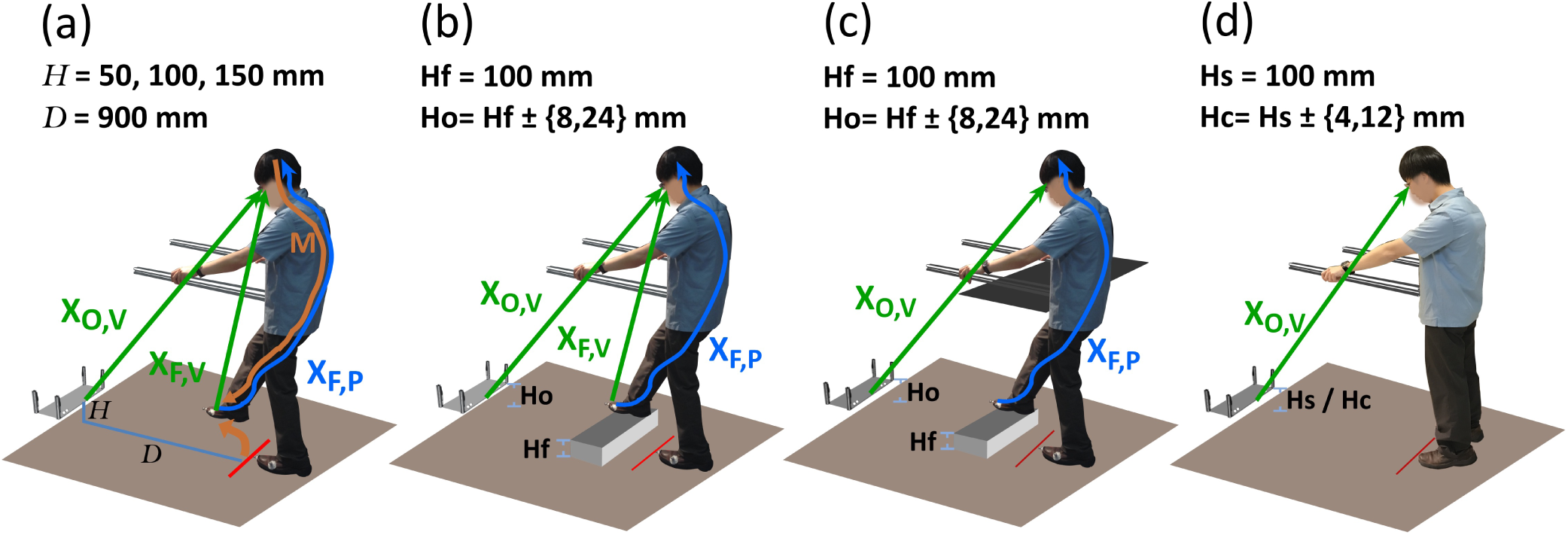
Experimental set-up. Participants stood inside an automated robotic platform, 900 mm away from the obstacle. They undertook four tasks. In **(a)** FOot HEight POsitioning (FOHEPO) task, participants lifted one of their feet to match the height *H* of the obstacle. Three target heights were examined (50 100 and 150 mm). The FOHEPO task measured sensorimotor variability. Sensory variability in the context of the FOHEPO task was then measured by using **(b)** foot-obstacle two-alternative forced choice (2AFC), leg-visible condition. In each trial, one of the participant’s feet was lifted by a 100 mm high platform (**Hf** 100 mm). The obstacle height **Ho** was randomly chosen from 4 levels: 100-24, 100-8, 100+8, and 100+24mm. Participants chose whether the lifted foot or the obstacle is higher in each trial. We then conducted two more sensory psychophysics **(c)** and **(d)** to compute sensory variability originated from individual sensory modalities, including vision and proprioception. In **(c)** Foot-obstacle 2AFC, leg-invisible condition. The procedure was identical to **(b)**, except that the visual information of foot height was blocked by a piece of A1 size black paper. **(d)** Obstacle 2AFC. Each trial consisted of two successive observation intervals. The standard stimulus (**Hs** 100 mm) randomly appeared in one of the two intervals. The comparison one **Hc** was randomly chosen from 4 levels: 100-12, 100-4, 100+4, and 100+12 mm. Each interval presented the stimulus for 2 seconds. Between these two intervals, actuators moved the obstacle to the position of the second interval. Participants then chose the interval containing the higher height between the two at the end of each trial

**Figure 2.**
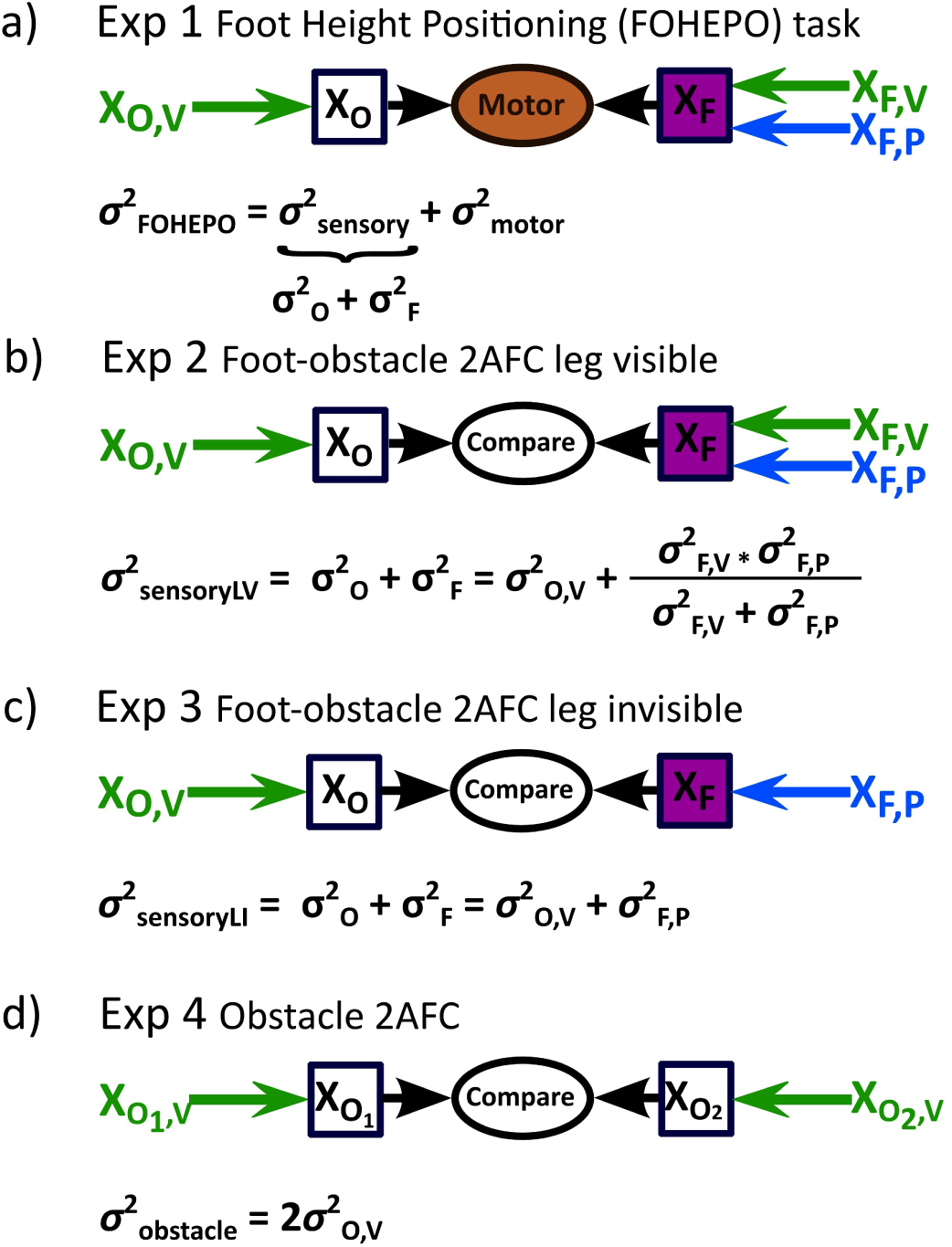
Models to represent and compute the variability of each task **(a)** In the FOHEPO task, task variability, i.e. sensorimotor variability, is the sum of sensory and motor variability^48^. The motor policy formed for the FOHEPO task is based on all available sensory information, including sensory estimates of the height of the obstacles (*X*_*O*_) and sensory estimates of the height of the own foot (*X*_*F*_). Therefore, the sensory variability of the FOHEPO task is the sum of foot height estimate variability 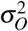 and obstacle height estimate variability 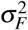 **(b)** We directly measured the sensory variability in the foot-obstacle 2AFC leg-visible condition. As has been mentioned, the sensory variability, expressed as the variance of encoded position estimates of foot and obstacle 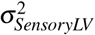, is the sum of of the variances of obstacle 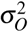 and the foot height estimates 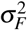^71^. Sensory estimation of the obstacle height is represented by the visual information. Therefore, the sensory variability of obstacle height 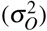 equals to the visual variability of the obstacle height 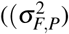. Sensory estimates of foot height in the leg-visible condition are formed by combining the variance of proprioception 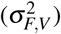 and vision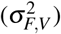 in a Bayesian optimal way. **(c)** When the lifted leg is invisible, foot height is estimated only based on proprioceptive representation and thus 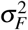 equals to 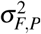. Total sensory variance 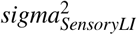 is still the linear sum of 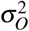 and 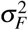. c.) In the obstacle 2AFC, the visual estimates of obstacle positions at two intervals of a trial are compared. We assumed the variances of obstacle height estimates remain constant in two intervals. From signal detection theory of psychophysics^72^, we know 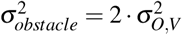, where 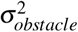 is attainable from psychometric function of obstacle 2AFC responses.

**Figure 3.**
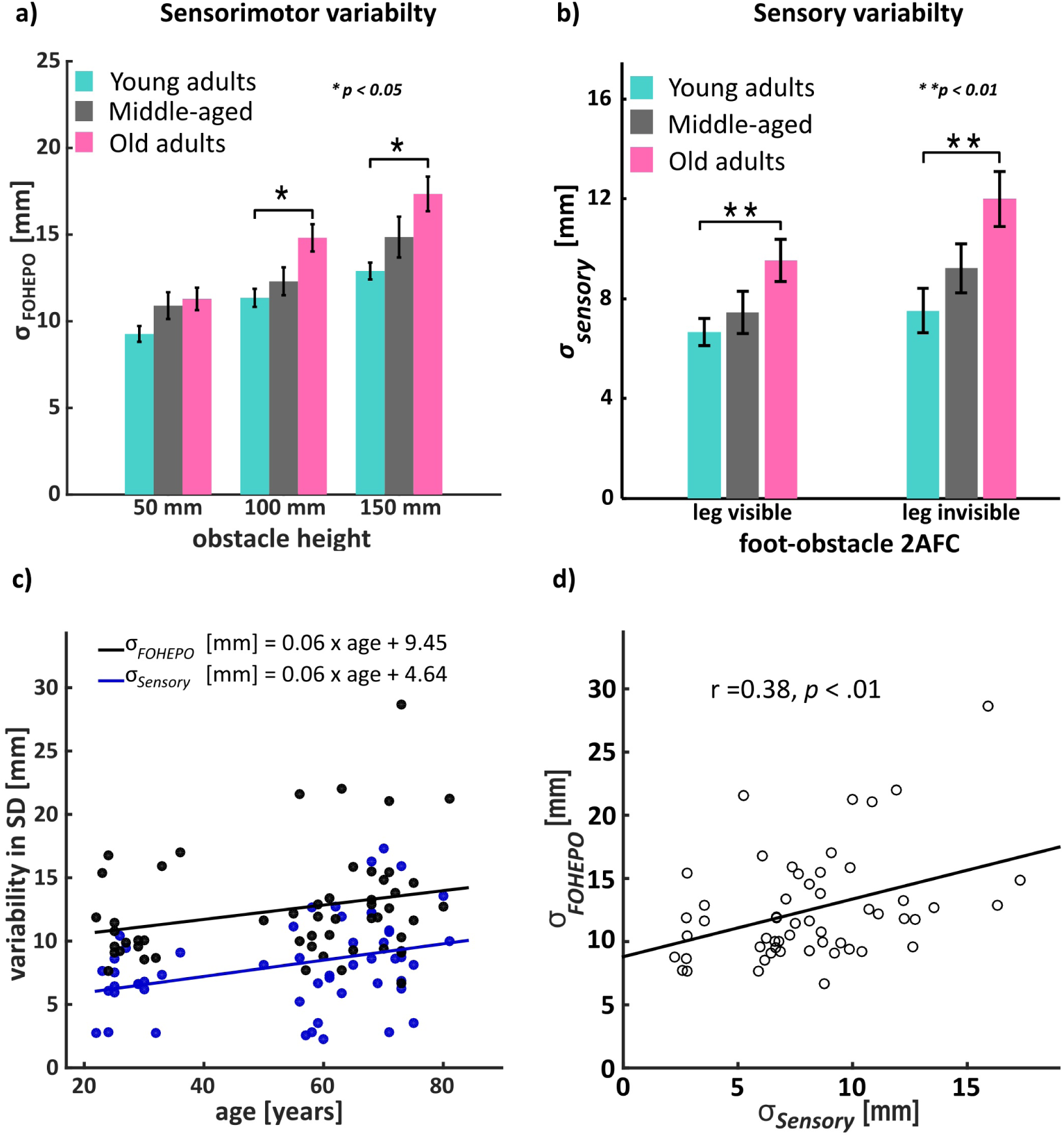
Sensorimotor and sensory variability under different experimental conditions. **(a)** Group means of sensorimotor variability of the FOHEPO task of each height condition. Significant age group differences were found in 100 mm (*p* = .012) and 150 mm (*p* = .014) conditions. **(b)** Group means of sensory variability in the foot-obstacle 2AFC, leg-visible and-invisible conditions. Elderly subjects had higher levels of sensory variability than the young subjects in both leg-visible (*p* = .032) and leg-invisible (*p* = .015) conditions. **(c)** Scatter plot of age and both sensorimotor and sensory variability in the 100 mm condition as a linear function of age. Age was a significant, positive predictor of both sensorimotor (*r* = .30, *p* = .026) and sensory variability (*r* = 0.35, *p <* 0.01). Across adulthood, sensorimotor and sensory variability both increased at 0.06 mm per year averagely. Similar graphics showing the sensorimotor variability of the 50 mm and 150 mm conditions versus age may be found in the supplementary material figure 2. **(d)** There was also a significant positive correlation between sensorimotor and sensory variability (*r* = .38, *p* = .004). The results of **(c)** and **(d)** imply that the sensory noise being one cause underlying more variable movement in ageing.

### Experiment 2 & 3: Sensory variability in the foot-obstacle two-alternative forced choice (2AFC)

To quantify sensory variability in the context of the FOHEPO task, we used foot-object height discrimination psychophysics. Sensory variability (Fig. 3 b.) showed significant age group (*F*_(2,52)_ = 6.07, *p* = .004) and condition (leg-visible versus leg-invisible) (*F*_(1,52)_ = 8.40, *p* = .005) effects, but no interaction. Pair-wise comparison found that elderly had higher levels of sensory variability than the other two groups in both leg-visible (*p* = .032) and leg-invisible (*p* = .015) conditions. This means that total sensory variability levels in elderly adults were significantly greater than those in young adults. Moreover, when comparing differences between leg-visible and-invisible conditions(mean*±* 95% confidence intervals), the increased amount of sensory variability in elderly adults were greater (leg-visible *vs* leg-invisible: 9.5 *±* 4.2*mm vs* 12.0 *±* 5.1*mm*; an increase by 26%) than young adults (leg-visible *vs* leg-invisible: 6.7 *±* 2.2*mm vs* 7.5 *±* 3.7*mm*; an increase by 12%). This implied that absent visual information of foot height negatively affected discrimination performance of all participants but the deterioration was more remarkable among elderly than young. When directly examining the differences of variance between the two conditions 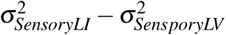, we discovered that elderly adults comprised significantly higher variance in the leg-invisible than in the leg-visible condition (*p* = .018, one-tailed t-test). While in young and middle-aged, the variance in two conditions were not statistically different (young *p* = .12; middle-aged *p* = .06).

We performed a linear regression on sensory variability against age. As can be seen from Fig. 3 c. (blue dots and line), age was a significant predictor of sensory variability. The regression equation was sensory variability (mm) = 4.64 (mm) + 0.06 *⋅* age (*r* = .35, *p* < .01). We then tested the hypothesis that raised sensory variability results in raised sensorimotor variability using correlation statistics. There was also a positive correlation between sensorimotor and sensory variability (Fig. 3 d.; *r* = .38, *p* = .004). A partial correlation between an individual’s sensorimotor variability at 100 mm and age whilst controlling the effect of sensory variability was also conducted. We found no positive partial correlation between sensorimotor variability and age whilst taking account the mediating effect of sensory variability (*ρ* = .14, *p* = .31). This result indicates that sensory variability had a significant influence on ageing-related sensorimotor variability increases.

#### Sensory variability decomposition

We also applied sensory psychophysics of object height discrimination and used the psychometric parameters acquired from all three sensory psychophysics to compute sensory variability of both vision and proprioception (see methods).

##### Proprioceptive variability of the foot estimates

As can be seen from Fig. 4 a., there was a significant age group effect (*F*_(2,44)_; *p* = .048). Pairwise comparison showed that elderly adults had higher proprioceptive variability compared to young (Bonferroni *p* = 0.055; LSD *p* = .018). The middle-aged group did not have significant differences to any of the other two groups. A linear regression on proprioceptive variability against age showed a significant positive correlation between proprioceptive variability and age (Fig. 4 b.; *r* = .30; *p* = .048). There was also significant positive correlation between sensory and proprioceptive variability (supplementary material figure 3.a.; *r* = .53; *p* < .001). Importantly, the result of a partial correlation between sensory variability and age whilst controlling the effect of proprioceptive variability was not significant (*ρ* = .20, *p* = .19), suggesting that the ageing-related increase of sensory variability likely due to the increase proprioceptive variability. It should be noted that we excluded some individual’s data (4 out of 17 in young, 4 out of 17 in middle-aged and 2 out of 21 in older) from statistical analysis because negative values of 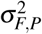 were obtained when using eq. (3) and (4) to extract proprioceptive variability.

**Figure 4.**
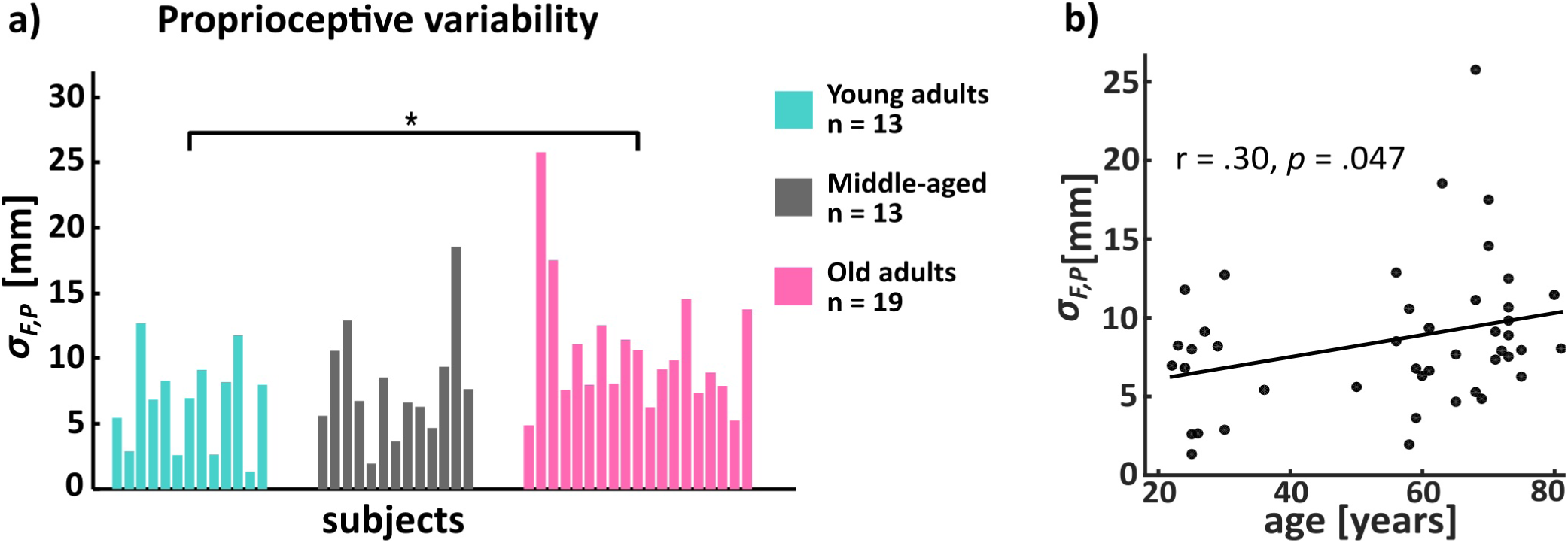
Proprioceptive variability **(a)** There was a significant age group effect (*F*_(2,44)_; *p* = .048). Pairwise comparison showed that elderly adults had higher proprioceptive variability compared to young (Bonferroni *p* = 0.055; LSD *p* = .018). Each bar represents single subject. It should be noted that the sampling numbers were different from those of sensory experiments. Some participants’ data (4 out of 17 young adults, 4 out of 17 middle-aged and 2 out of 21 old adults) had to be excluded because negative values of 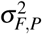 were obtained when using eq. (3) and (4) to extract proprioceptive variability. **(b)** Scatter plot and of age and proprioceptive variability. Age was a positive predictor of proprioceptive variability (*r* = .30; *p* = .048).

##### Visual variability of the obstacle and foot estimates

The visual variability of obstacle height had neither age group difference (*F*_(2,58)_ = 1.3; *p* = .28) (Fig. 5 a.) nor correlation (*p* = .20) with participants’ age. Additionally, the result of a partial correlation between sensory variability and age whilst controlling the effect of visual variability remained to be significant (*ρ* = .32, *p* = .033), indicating that visual variability had very little influence in controlling for the relationship between sensory variability and age. However, we also found that only in the elderly group, there were significant positive correlations between the visual variability of obstacle height and sensorimotor variability (Fig. 5 c.; *r* = .47; *p* = .033) as well as sensory variability (*r* = .49; *p* = .023, see supplementary material figure 3.b.). This implied that individual differences of visual precision affect motor precision of the elderly group but not the other two groups.

**Figure 5.**
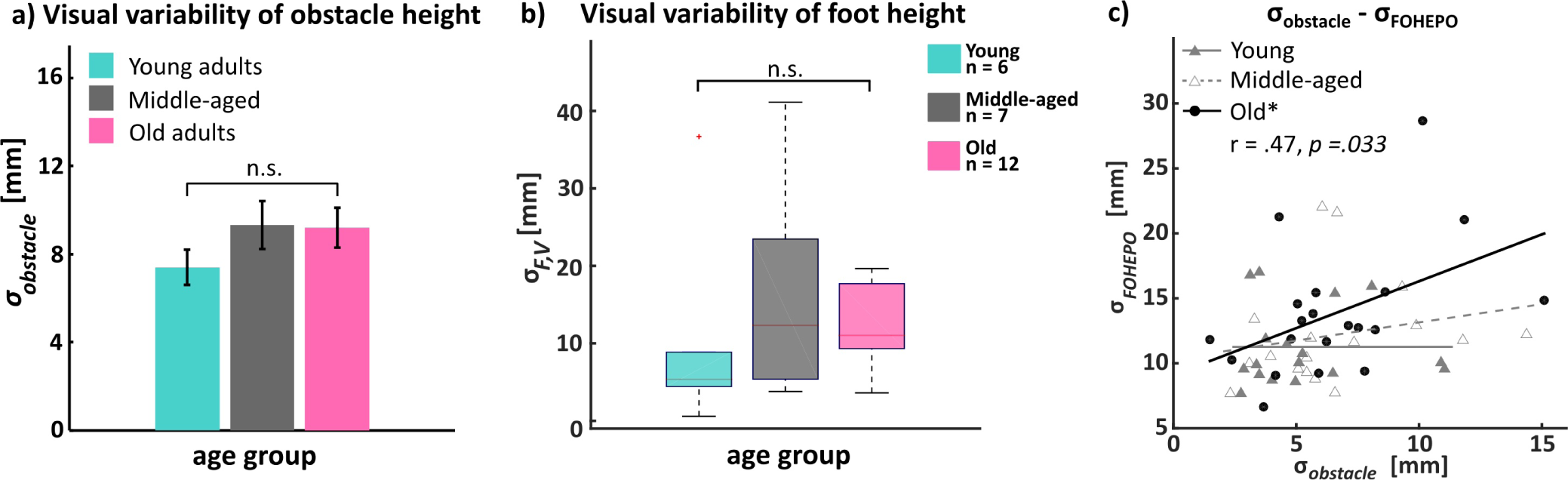
Visual variability of obstacle and foot height **(a)** mean visual variability of obstacle height *σ*_*obstacle*_ of each age group. There was no significant age group differences by ANOVA. Error bars indicate SEM. **(b)** Visual variability of foot height *σ*_*F,V*_. Individual visual variability was acquired by using eq.(2), which assumed people followed Bayesian rule when integrating visual and proprioceptive information to estimate their foot heights. There was no significant age group effect. **(c)** Scatter plot of visual variability of obstacle height and sensorimotor variability. Only in the elderly group, there was a significant positive correlation between visual variability and sensorimotor variability *r* = .47; *p* = .033. In the young and middle-aged groups, no such relationship was observed.

Based on the assumption of optimal integration rule, we also attempted to extract visual variability of foot estimates using eq. (2) (see Method section for details). However, negative values of 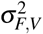 were found in a notable proportion of participants. After excluding those cases, the medium of 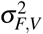 was found to be 5.4mm in the young (N=6), 12.3mm in the middle-aged (N=7) and 11.0mm in the older (N=12) (Fig. 5 b.). The result of Kruskal-Wallis H test was not significant (*p* = .24).

#### Modelling tripping-over occurrence by age

Using our sensorimotor variability data, we built a probability model to predict tripping-over numbers on staircases at different ages and compared with community-based statistics. As can be seen from Fig. 6 a., we predicted that the probability of tripping-over on a 150 mm staircase increased from less than 10^*−*^7 per step in people’s 20s to 10^*−*^4 per step in their 80s. Accordingly, expected tripping-over numbers over a period of 2 years increased from 5 *×* 10^*−*^3 to 3 (Fig. 6 b.). Compared to real-life fall-upon-stair data from the population^36^, the trend of the increases by age was generally in line with each other. Fall numbers were around 3-10 times lower than predicted tripping-over numbers. This is reasonable because a tripping-over does not always result in a fall event. However, as can be noted, the trip-over/fall ratio increased with age.

**Figure 6.**
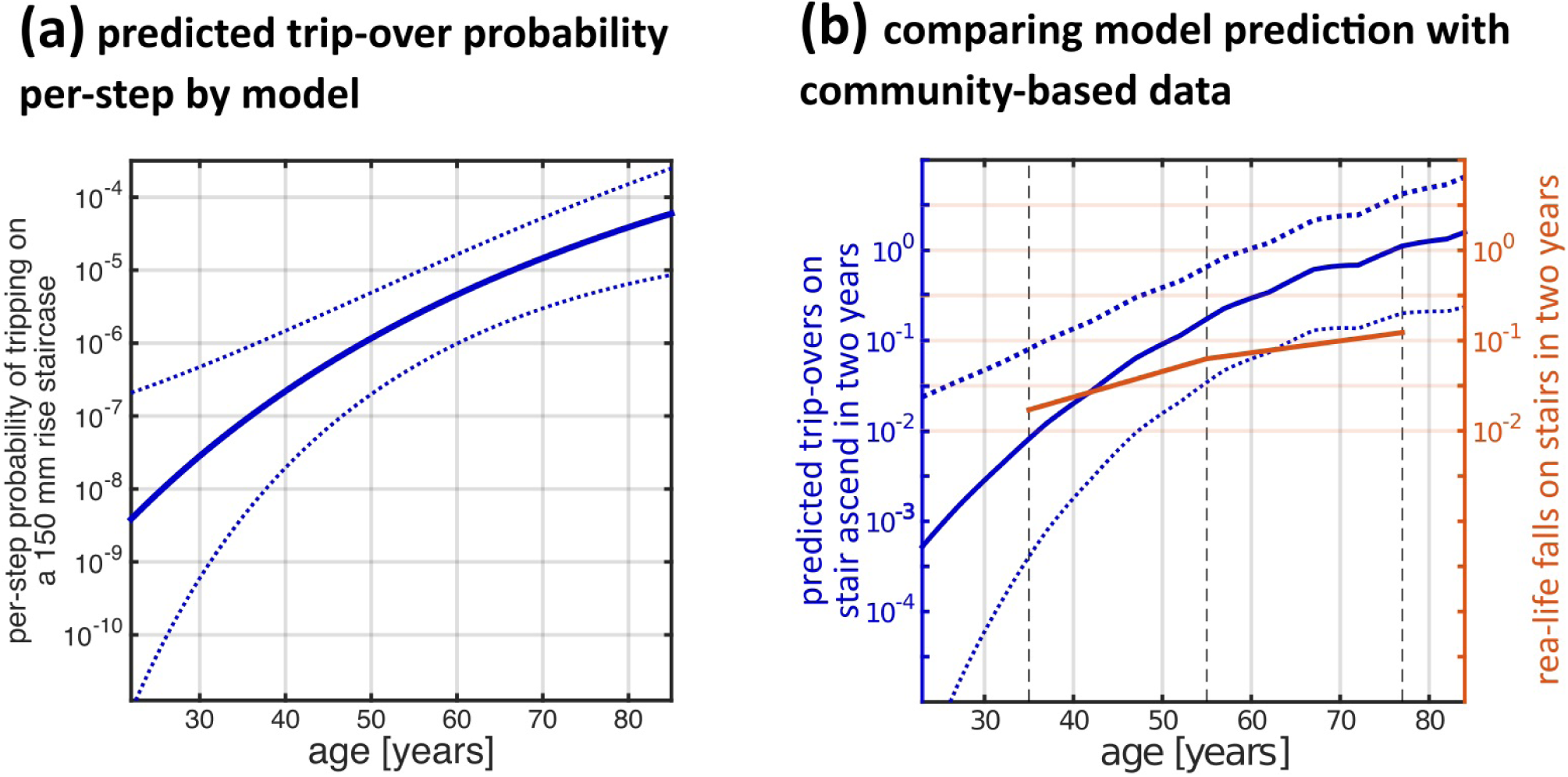
**(a)** Tripping probability modelling as a function of age for healthy adults aged 22-84 years during stepping onto a 150 mm rise staircase. The model was based on the linear function of sensorimotor variability of the FOHEPO task, 150 mm height condition, by age. Tripping probability increased around 10^3^ times from 20+ to 80+ years. Dashed lines represent the 95% confidence intervals. **(b)** Compare model prediction to reported data. The blue line represents two year expected trip-over numbers by age using the model shown in **(a)**) with adjustments of activity levels according to Health Survey for England 2008^77^. Here we assumed that a 50 year old person climbs 10 flights of 15 cm rise stairs daily^76^. As can be seen, trip-over occurrence increased nearly monotonically, from less than 10^*−*2^ at 22 years to about 3 times at 84 years in a two year period. A brief and small decline of expected trip-over number in the 70s was due to decreased activity levels and thus total stair climbing numbers. The red line represents community-based fall numbers of three age groups: young (mean 35 years), middle-aged (mean 55 years) and elderly (mean 77 years) people, in a two year retrospective study done by Talbot et al.^36^. As can be seen, the trend of model prediction was broadly similar to community-based data. Dashed lines represent the 95% confidence intervals.

## Discussion

Greater gait variability, specifically, greater MFC variability, has been identified as an important marker of increased fall risk in the elderly, yet the neural computations underlying changes of behavioural performance with age has mostly been unexplored. Here, we used a novel sensorimotor task with the shared biomechanics of obstacle negotiation: the FOHEPO task, to measure the sensorimotor variability of foot placement. We demonstrated that sensorimotor variability increases in the elderly. We then used sensory psychophysics to quantify sensory variability in the context of the FOHEPO task and showed it higher in the elderly. Moreover, while both sensorimotor and sensory variability positively correlated with subjects’ age, the correlation between sensorimotor variability and age was explained away in a partial correlation whilst controlling sensory variability. This suggests increased sensory variability responsible for the age group differences in sensorimotor variability of the FOHEPO task. We further dissected quantitatively the three key components of sensory variability: i.e. proprioceptive variability of foot estimates, visual variability of foot estimates and visual variability of obstacle estimates, by conducting two more sensory psychophysics and applying the computational model of sensorimotor integration. Our key finding is that proprioceptive but not visual variability increases among elderly participants. Finally, we applied computational models of sensorimotor control parametrised by our sensorimotor variability results to predict how age-dependent changes increase falls triggered by trip-overs in the elderly.

The two common ageing motor features are increased end-point variability and slower movements^16–22^. In this study, we focused on end-point variability and provided sufficient time for task completion. We found elderly participants spent 13% more time on foot matching (averagely 1.69 seconds in the young while 1.91 seconds in the elderly). (Supplementary Table 1, column 2). This result found in the lower limb is in agreement with the observations in perception-action tasks in the upper limb^16, 20^. Additionally, it is consistent with the work of biomechanics of ageing, where older adults display higher gait variability, independent of slower walking speed^37, 38^. Moreover, with abundant time to respond (5 seconds to observe and prepare and 4 seconds to move, 99.4-99.8% trials completed well within the allowed trial time), it is rational to conclude that the increased sensorimotor variability with age did not result from elderly trading foot placement accuracy with speed. In fact, this conclusion was supported by a finding that no speed-accuracy correlation was shown in the foot placement data (supplementary material figure 5).

We used regression to examine how sensorimotor variability changes over the course of adulthood and found an increase roughly by 0.07 mm per year. The average sensorimotor variability of young people was 11.2 mm, so, accumulatively, we predicted an increase in sensorimotor variability by 29% from 27 years, the mean age of our young group, to 73 years, the mean age of our elderly group. This trend appears to be significantly linear and not characterised by sudden non-linear jumps, as evidenced by comparing the goodness of fit to non-linear (quadratic) functions. Indeed, previous studies identified gait variability increases as early as the age of 50^39, 40^. Variability in controlled perception-action tasks across adult lifespan has been less investigated. Two reports observed that the grip force steadiness remained stable until age 60^42, 43^. The discrepancies between our finding and those in the force studies^31, 43^ could have been caused by differences in task nature. Our FOHEPO task needs integration of multiple sites in the sensorimotor system, from sensory organs, primary sensory areas, multisensory cortices, cortical and subcortical motor areas, spinomotor neurons to muscles. Studies have shown that different regions of the sensorimotor system undergo dissimilar trajectories along the ageing course^19^. For example, there appears to be no evidence of spinal motor neurone loss before the age 60^44, 45^ while in the brain, the cortical thickness^46^ and neuron number^47^ of motor areas decrease continuously across adulthood. The distributed computations needed for the FOHEPO task may be the reason that the performance degraded continuously but also more gracefully across age. Understanding the effect of age on sensorimotor variability across adulthood helps us to recognise early ageing related changes of the sensorimotor system, in which potential clinical targets for early detection and activity modifications to prevent later impairments lay.

Sensorimotor variability is composed of sensory variability, motor variability and internal mechanisms mediating the two^48^. We find that sensory variability, measured via the foot-obstacle 2AFC task in the leg-visible condition (i.e. experiment 2), was significantly higher (by 42%, 2.8 mm) in the elderly than the young. The quantitative increase of sensory variability matched which in sensorimotor variability (by 30%, 3.4 mm). More importantly, when controlling for the effect of sensory variability by partial correlation, the positive correlation between sensorimotor variability and age was explained away. We concluded that sensory variability is the primary driver for age-related increases in sensorimotor variability. Our quantitative result was consistent with qualitative findings of anatomical^49, 50^, physiological^51^ and behavioural^52^ data which reflect decreased sensory functions in the elderly.

Sensory variability in our task constituted primarily three factors, visual variability in estimating the obstacle height, visual variability in estimating the own foot heights and last but not least, proprioceptive variability in estimating the own foot height. We were able to demonstrate, by combining our assay of sensory psychophysics experiments, that proprioceptive variability increased significantly by 56% in the elderly group. Furthermore, the ageing-related increases of sensory variability were explained away by proprioceptive but not visual variability, as demonstrated by the results of partial correlations. This provides evidence supporting the importance of proprioception to gait control. Previous studies revealed that impaired proprioception during ageing impacts stance and balance control^50, 53^ but they focused on static scenarios, not dynamic movements of the feet. We were able to show here that increased proprioceptive variability affects critically sensorimotor variability and as a result, the variability with which an ageing person clears over an obstacle (foot clearance variability).

As visual performance loss and attributed sensory variability are commonly assumed to be a prominent characteristic during ageing, an absence of raised visual variability in our elderly participants was surprising. Ageing-related visual dysfunction can be categorised into optical and neural changes. It has been shown previously that the eye’s optical degradation due to increased light scatters by the clouding of aged lenses, i.e. cataract, is a primary source of increased visual variability during ageing^49^. We evaluated our elderly participants’ health status questionnaires, which showed that they had corrected normal sight and also a very low (2 out of 26 in older group; 1 out of 18 in middle-aged) prevalence of cataracts compared to 40% incidence in population-based studies^54, 55^. Thus, we can conclude that the visual system in our study population was little affected by age and therefore could not explain the increase of sensorimotor variability observed in the sampling population. This underlines the importance of our finding that proprioceptive variability the key cause of sensorimotor variability increases, as it provides the theoretical foundation of a rehabilitation pathway: previous work has shown that supervised physical training of proprioceptive capabilities promotes retention^56^ and even improvements on proprioceptive performance^57^.

When the visual information of foot height was blocked, foot-obstacle height discriminability declined by 12-26%. This shows that subjects did integrate foot height estimates from both vision and proprioception to boost sensory representation precision, as would be expected. However, the degree of precision raised by visual information differed among age groups. There was no significant improvement in the young and middle-aged. In contrast, the elderly group benefited substantially from using vision. This result is in-line with research discovering that the elderly compared to the young rely more on vision to maintain balance^58^. It is also compatible with previous studies showing enhanced benefits from multisensory integration in the elderly compared to the young^59^. It should be noted that the so-called ‘enhanced benefits’ does not imply elderly have a more efficient multi-sensory combination mechanism, but indicates that they experience greater benefits from the multisensory combination as unisensory estimates degrade. Shifting weighting on the remaining intact sensory modalities/modality is an effective compensatory mechanism but also raises concerns. Research has repeatedly shown, elderly people are less flexible and efficient when there is an acute change in sensory information reliability that they have to reweigh to regain “optimal” integration^58, 60, 61^. This means when the relied modalities/modality of information suddenly deteriorate(s) or disappear(s) in some circumstances (such as rapid changes of illumination, which commonly occur in cinemas), substantial increases in erroneous sensory estimates and thus motor outputs are more likely to happen. This assumption is supported by our finding that a positive correlation between sensorimotor variability and the visual variability of object height only existed in the elderly, but not the young and middle-aged. The result reflected the raised influence of visual uncertainty on overall sensorimotor variability with age. From an application perspective, this discovery opens up a new direction for ageing friendly space design^62^. It also indicates the importance of developing training activities enhancing sensory reweighing policy in the elderly.

The FOHEPO task can potentially be used a fall risk screening tool. To evaluate the feasibility, we built up a pipeline to compute expected biyearly trip-over numbers on stair ascend based our derived mathematical models of age-related changes in variability. The expected trip-over numbers increased around 10^3^ times, from around 10^*−*3^ at 22 years to approximately 1 to 2 at 85 years. We also compared the model prediction to a two-year cohort-based study on fall counts upon stair climbing^36^. Similar to the model prediction, the number of falls on stairs increased with age and were always lower than the modelled trip-over numbers. This is reasonable considering that not every trip-over eventually results in a fall. However, there were also discrepancies. While trip-over numbers in the model increased by around 50 times from 35 to late 70s, fall incidence in real life raised by only seven times. The ratios between trip-over events and falls were thus 2.75 at 35 years and 15.60 at 77 years. One possible explanation is that falls on stairs in the young may be more likely triggered by causes other than increased movement variability, such as risky activities on stairs. Moreover, our model assumed that the mean clearance remains constant along the adulthood. However, there has been research suggesting that older adults adapted to a more conservative pattern with a higher degree of joint flexion during obstacle negotiation, which could lead to increased mean stair clearance (see review^63^).

As with all lab-based tasks, the question remains, how well the results in the controlled FOHEPO task can be generalised in real-life situation. Our lab-based results should be validated in a larger ageing cohort using appropriate tools that enable us to associate intra-individual sensorimotor variability with foot clearance variability. One fundamental difference between the FOHEPO task and daily walking is that subjects in our experiments could pay full attention to the task while distractions from the environment complicate real-life navigation. Research has revealed that MFC impairment with age becomes more noticeable as their cognitive control resources were diverted to other tasks^64^. Therefore, the results of our controlled task show the robustness of age-related sensorimotor variability differences and it is reasonable to assume that this age group differences will maintain significant in walking scenarios. Further, future test populations should include neurodegenerative patients as studies have shown that they also present with increased gait variability^65, 66^.

To conclude, the current study, to the knowledge of the authors, is the first experimental evidence to show a connection between increased sensorimotor variability and increased foot clearance variability. Importantly, we systematically examined the relationship between changes in individual sensory modalities, overall sensory variability and sensorimotor variability. We demonstrate that the experimental protocols can pinpoint a specific deficit of the sensory systems, namely increased proprioceptive variability, critically linking to increased sensorimotor variability of the tested ageing subjects. The finding provides the neural control understanding of falls and also has practical implications for the future development of personalised risk screening tools and targeted rehabilitation exercise.

## Methods

Four experiments were conducted (Fig. 1). In experiment 1, the FOot HEight POsitioning (FOHEPO) task, we measured the sensorimotor variability of foot stepping. In experiment 2, we used height discrimination between the foot and obstacle to quantify sensory variability of the FOHEPO task. The procedure of Experiment 3 was identical to experiment 2 except that the visual information of the foot height was not available for height discrimination. In experiment 4, we applied obstacle height discrimination to examine the visual variability of obstacle heights. The purpose of experiment 3 and 4 was to compute sensory variability of individual sensory modalities. We then built a probability model to predict fall occurrence as a function of age based on the data from experiment 1.

### Participants

Seventy-four people, including 30 young (13 females; mean 27.0 (S.D. 3.9) years, pre-defined range 20-40 years), 26 elderly (11 females; mean 73.3 (S.D. 4.4) years, pre-defined range 66 years and beyond) and 18 middle-aged (10 females; mean 59.5 (S.D. 3.9) years, pre-defined range 45-65 years) volunteered in the FOHEPO task. Young participants were graduates of Imperial College. Participants 45 years or beyond were volunteers from the local community or Imperial College. All subjects gave informed consent according to university protocols and completed a health questionnaire to ensure they had no past or present conditions that could interfere with the sensorimotor control and Waterloo Footedness Questionnaire-Revised (WFQ-R)^67^ to determine their dominant foot. All but one young participant were right dominant. Out of 74 who completed the FOHEPO task, 17 young (9 females, mean age 27.5 (S.D. 4.0) years), 22 elderly (11 females, mean age 69.8 (S.D. 5.8) years) and 17 middle-aged (9 females, mean age 59.5 (S.D. 3.9) years) volunteers completed the foot-obstacle 2AFC. Eighteen young (10 females; mean 28.6 (S.D. 4.7) years), 23 elderly (11 female, mean age 69.8 (S.D. 5.9) years) and 17 middle-aged (9 female, mean age 59.5 (S.D. 3.9) years) completed the obstacle 2AFC task. The reason of that some participants left sensory experiments undone was relocation, except that one elderly participant withdrew after completing the FOHEPO task and foot-obstacle leg visible 2AFC because of an exercise injury, and one middle-aged participant lost contact. Data was unusable in one of the FOHEPO task (in the young group) and one of the foot-obstacle 2AFC (in the elderly group). There were no significant group differences in mean height or weight or gender distribution.

### Stimuli and apparatus

All the experiments were performed in the FOHEPO workstation, a closed-loop robotic environment we built for high precision motor and sensory psychophysics on the lower extremities^68^. The workstation constituted of a supportive metal frame with its rear side open for a participant standing inside the workstation. The stimulus (We called it the ‘obstacle’ because it served as an obstacle that a subject had to step to.), a white polystyrene sheet L 760 mm *×* W 300 mm *×* H 5 mm was located at the front side of the workstation and surrounded by black workstation walls and floor. Four Firgelli linear actuators (Firgelli L16-140-63-12-S, Firgelli, Victoria(BC/Canada) drove the obstacle moving vertically from 0 (= ground level) to 150 mm. Three Optitrack Flex-13 cameras (Natural Point, OR, USA) on wall shelves above the front margin of the workstation continuously tracked the positions of participants’ feet and the obstacle.

### Procedure

#### Experiment 1: FOHEPO task

(Fig. 1 a.) Participants stood 900 mm from the obstacle while holding safety handrails. Each trial started with actuators moving the obstacle to the target height. Five hundred ms after the obstacle reached its target level, a “countdown” phase with an audio signal being “three-two-one-beep” was delivered. Subjects got prepared but kept still when hearing “three-two-one” and started matching the designated foot to the obstacle on the ‘beep’.They kept the moving foot parallel to the ground and flexed their knee and hip to raise the foot. When they considered their foot matching the obstacle height, they pressed a button on the left handrail to transmit a flag for time monitoring purposes to the PC-controlled MATLAB Module. Subjects kept their foot at the same level for 1000 ms until they heard another audio signal, which indicated the end of a trial and also provided performance feedback using real-time height difference extracted from motion tracking data (positive feedback when the difference between the foot and the obstacle less than 35 mm; otherwise negative feedback). If a participant did not press the button within 4000 ms after a “beep”, the trial was terminated automatically and labelled as “mistrial”. Participants were informed before the experiment that they would have one more chance to repeat mistrials at the end of each block and that the speed of reaction was not the primary measure on their behaviours and they could spend as much time on a trial provided that they complete a trial within 4000 msIf a participant failed to complete a trial in the second try, that trial was excluded from data analysis. Three target heights (50, 100, and 150 mm) were presented with 20 trials for each foot, resulting in a total of 120 trials.

#### Experiment 2 & 3: Foot-obstacle two-alternative forced choice (2AFC) – leg visible and invisible

(Fig. 1 b. and c.) A constant-stimulus method with a 2AFC task was used. Each trial consisted of the simultaneous presentation of one of the participant’s feet, which was lifted to 100 mm above the ground by a platform, and the obstacle. Foot heights were taken as standard stimuli. The obstacle heights, varying from *±*8 and *±*24 mm to the standard were presented as comparison stimuli. The participant chose the higher stimuli by pressing buttons on a numeric keyboard, on which button “F” meant “foot” higher and button “O” meant “obstacle” higher. Two conditions, one with the lifted leg visible and the other with the leg visually blocked, took place on two separate days. Each condition contained 10 blocks of 12 trials with both feet being tested. The orders of the two conditions and both feet were counterbalanced among participants.

#### Experiment 4: Obstacle 2AFC

(Fig. 1 d.) A constant-stimulus method with a 2AFC task was used. Each trial consisted of the sequential manifestation of two obstacle stimuli. The height of the standard was 100 mm. The comparison was a set of heights from *±*4 and *±*12 mm to the standard. Between trials and intervals, linear actuators moved the obstacle. Participants closed their eyes between intervals and trials. Sound signals indicating the beginning and end of an interval were played to instruct participants when to observe the height of the obstacle front border. At the end of a trial, participants chose the higher interval by pressing buttons on a numeric keypad, on which button “1” represented the first interval and button “2” represented the second. There were 10 blocks of 12 trials in one experiment.

### Data processing and statistical analysis

Data analysis was performed by MATLAB (Mathworks, Inc. Natick, MA, USA). In the FOHEPO task, the average of 1 second (120 frames) front marker height data, beginning from button press time, was regarded to be foot height, so as the height of the obstacle. Statistical analysis was performed by SPSS (IBM Corp. Released 2013. IBM SPSS Statistics for Windows, Version 22.0 Armonk, NY: IBM Corp.). The Kolmogorov–Smirnoff test was used to check the normality of the data. For normally distributed data sets, we used t-tests (two-tailed, unless otherwise stated) and mixed-design ANOVAs. Bonferroni’s method was used for pairwise comparisons unless otherwise stated. For data sets that were not normally distributed, the nonparametric Kruskal-Wallis H test was used. Outliers were detected by Tukey’s method^69^, taking (Quartile3) + (3 IQR) as a cut-off value.

#### Fitting psychometric functions and computing sensory variability

The experiment 2, 3, & 4 involved estimating sensory variability using psychometric function parameters. We performed computations as follows:

Raw responses were the proportion of trials in which a comparison stimulus was judged higher than a standard at each comparison level. We used raw responses to fit psychometric functions using a cumulative Gaussian distribution^70^.

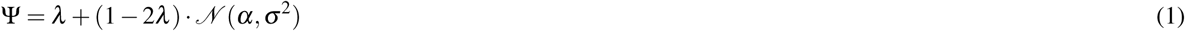

Performing the curve fitting enabled us to obtain *σ* ^2^ the variance of sensory estimate distribution of each sensory experiment^71, 72^. Other terms in the equation include Ψ the probability of a comparison higher response given a specific height difference, *α* the comparison height corresponding to the 50% point of the function and *λ* the lapse rate. In experiment 2 & 3, *σ*^2^ is the linear sum of obstacle estimate variance and foot estimate variance and can be expressed as the following equations, where *σ* ^2^ was the left-sided term of each equation. (also see Fig 2)

Experiment 2: foot-obstacle 2AFC – leg-visible(LV)

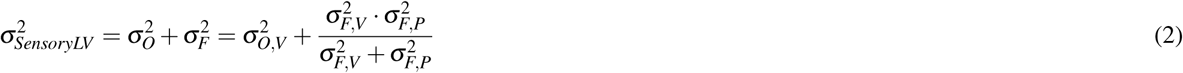

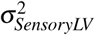 represents the variance of the difference distribution and also the sensory variability 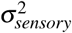 of the FOHEPO task. The variance of foot estimates in this experiment is a product of visual variance of foot estimates and proprioceptive variance of foot estimates, where the overall foot estimate variance equals 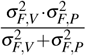 by assuming foot height estimation follows the Bayesian integration rule^73^.

Experiment 3: foot-obstacle 2AFC – leg-invisible(LI):

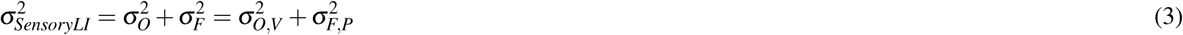

With only proprioceptive information of foot positions available, 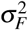 in experiment 3 thus equals to 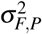.

In experiment 4 (Obstacle 2AFC),there is such a relationship between the measured 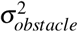 and the variance of obstacle estimates 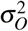 based on literature^72^:

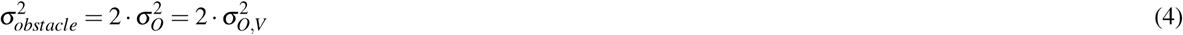

According to eq. (4), 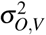 equals 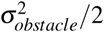. With 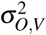 now being known, we could acquire 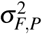 by subtracting 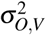 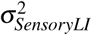 from 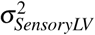 (based on eq. (3)). Last, since 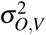 are measurable and 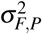 can be deduced, we can compute 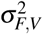 using eq. (2).

#### Modelling tripping-over occurrence

We computed yearly tripping-over occurrence as a function of age and then related to population data on falls during stair negotiation at different ages. To link falls per year to the trip-over probability per staircase we set up the following inference pipeline.

1. Trip probability per-step prediction: We built a model predicting the trip-over probability per step taken on negotiating structured obstacles, specifically ascending stairs with 150 mm risers. We fitted the sensorimotor variability data of the 150 mm condition of the FOHEPO task to a linear model which predicted the variability as a function of age

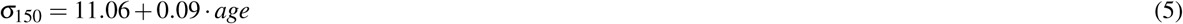 We assumed foot clearance data during stair climbing distribute normally based on literature^74^ and our own data (supplementary material figure 4). We used published foot clearance to low-rise staircases (as our 150 mm example case) from a study by Riener and colleagues^75^ as the mean *µ* and *sigma*_150_ of our model the standard deviation of foot distributions. We then computed the per-step probability of tripping-over *p*_*trip*_, i.e. the cumulative probability of foot placement lower than a 150 mm high staircase, using the following equation

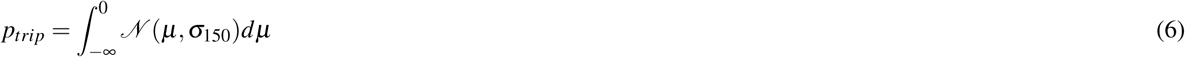
2. Expected trip-over numbers in a two-year period: We can calculate expected yearly trip numbers per age using the equation: tripping-overs per year = *p*_*trip*_ ⋅ stair climbs per year. As there were no population-based yearly stair climb counts available, we adapted data from a study conducted by Coupland and colleagues, which showed that people in their 50s in England climbed 10 flights of stairs daily on average^76^. We then normalised stair climbing counts according to activity levels (i.e. stair climbing hours per day) acquired from Health Survey for England 2008 by 10 year age groups (16-75+ years)^77^.
3. Comparing model prediction with survey data: We intended to compare trip-over numbers expected by the model at different ages to reported data. Only few studies documented both exact fall numbers and causes. A cohort study by Talbot and colleagues provided frequencies of total fall numbers, from 0 to 5+ falls, during a two-year period and activities prior to falling, in young, middle-aged and older groups^36^. We computed the expected two-year fall numbers while stair climbing of each age group *E*(fall as follows 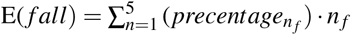, where *n*_*f*_ are the total number of falls.

## Acknowledgements

We express our sincere gratitude to the volunteers from (1) University of 3rd Age (2) Friends of Imperial College and (3) Imperial College London who kindly donated their time and energy to participate in this study. C.L. was supported by the Ministry of Education in Taiwan.

## Author contributions statement

C.L. designed the protocol, collected and analysed the data, implemented the probability modelling, wrote the manuscript and designed and prepared all figures, A.A.F. designed the protocol, analysed the data, and reviewed and edited the manuscript.

## Additional information

Competing financial interests: The authors declare no competing financial interests

